# Activity of spinal RORβ neurons is related to functional improvements following combination treatment after complete SCI

**DOI:** 10.1101/2024.03.01.583033

**Authors:** Nicholas J. Stachowski, Jaimena H. Wheel, Sebastian J. Atoche, Lihua Yao, D. Leonardo Garcia-Ramirez, Simon F. Giszter, Kimberly J. Dougherty

**Affiliations:** Marion Murray Spinal Cord Research Center, Department of Neurobiology and Anatomy, Drexel University College of Medicine, Philadelphia, PA 19129, USA

**Keywords:** locomotion, plasticity, spinal cord injury, afferent gating, epidural stimulation, BDNF, interneuron

## Abstract

Various strategies targeting spinal locomotor circuitry have been associated with functional improvements after spinal cord injury (SCI). However, the neuronal populations mediating beneficial effects remain largely unknown. In a mouse model of complete SCI, virally-delivered BDNF (AAV-BDNF) activates hindlimb stepping and causes hyperreflexia, whereas sub-motor threshold epidural stimulation (ES) reduced BDNF-induced hyperreflexia. Given their role in gating proprioceptive afferents and potential convergence point of BDNF and ES, we hypothesized that an enhanced excitability of inhibitory RORβ neurons would be associated with locomotor improvements. Ex vivo spinal slice recordings revealed that the excitability of RORβ neurons was decreased in mice with poor locomotor function after SCI, but was similar between the uninjured and ‘best stepping’ SCI groups. Further, chemogenetic excitation of RORβ neurons reduced BDNF-induced hyperreflexia and improved stepping, similar to ES. Our findings identify inhibitory RORβ neurons as a target population to limit hyperreflexia and enhance locomotor function after SCI.

## INTRODUCTION

Spinal circuits are capable of orchestrating a range of sensorimotor responses independent from the brain, including the patterned activation of lower motor neurons necessary for locomotion^1-3^. After complete spinal cord injury (SCI), locomotor circuits remain intact below most lesions^4-6^; however, lack of appropriate descending control of these circuits can result in the loss of function and/or the maladaptive gain of function (i.e. spasticity, chronic pain)^7-9^. Restoration of locomotor function after severe SCI will require the activation of silent spinal circuits and the reinstatement of a well-balanced control by spinal inhibitory mechanisms. Thus, combinatorial strategies capable of modulating various aspects of spinal sensorimotor circuitry will be necessary to compensate for the loss of descending control.

Increases in the expression of neurotrophic factors, particularly brain-derived neurotrophic factor (BDNF), have been associated with functional improvements following various strategies thought to engage locomotor circuits via sensory afferent pathways^10-14^. BDNF-TrkB signaling mediates neuronal plasticity^15,16^ and has roles in the development and maintenance of inhibitory neurotransmission^17-21^. Experimentally, viral delivery of BDNF (AAV-BDNF) is sufficient to promote stepping after spinal transection in adult cats^22^ and rats^23-25^. Continual overexpression of BDNF using AAV-BDNF is likely to have many actions, including increasing motor neuronal excitability and levels of both glutamatergic and GABAergic neurotransmission after chronic SCI^23-25^. Despite its effectiveness in improving stepping, viral BDNF treatment has also been shown to enhance afferent sprouting and often results in hyperreflexia and central sensitization in experimental models^23,24,26^, precluding the use of AAV-BDNF as a clinically relevant therapy. However, identification of the targets of BDNF mediating the beneficial actions would provide entry points to leverage in future therapies. Furthermore, how the BDNF-induced state of spinal excitability interacts with other potential therapeutics, and specifically epidural stimulation (ES), is poorly understood.

ES is a promising clinical intervention for recovery of motor function after SCI with demonstrated efficacy across numerous spinal systems^27,28^. ES is thought to modulate pre-motor and motor neuron excitability primarily through the recruitment of large diameter, myelinated afferents^29-31^. Initially developed and FDA-approved to treat intractable pain based on the classical gate control theory of pain^32-34^, clinical observations have since demonstrated that ES also reduces spasticity^35,36^, a prominent co-morbidity for SCI^37-39^. Despite the wide-ranging beneficial effects of ES, alone and as a supplement to motor rehabilitation after SCI^40-45^, remarkably little is known regarding the precise mechanisms of action^46^, particularly at the level of specific spinal neurons.

Spinal inhibitory interneurons control spinal excitability through both presynaptic and postsynaptic inhibition^47^. Interneurons involved in presynaptic inhibition of primary afferents can directly regulate sensory transmission of existing and sprouting fibers through primary afferent depolarization (PAD)^48-50^. Inhibitory interneurons in the medial deep dorsal horn which express the retinoid-related orphan nuclear receptor β (RORβ) presynaptically gate proprioceptive afferent input during locomotion^51^. Sensory-derived BDNF regulates the function of proprioceptive gating^20,52^ and modulates sensorimotor responses^53^. This is at least partly mediated by RORβ neurons which express TrkB receptors^51^. Moreover, genetic ablation of the RORβ population in the spinal cord, or selective removal of TrkB receptors from these neurons, results in a pronounced hyperreflexive gait^51^. Thus, RORβ neurons have the potential to be a convergence point mediating one of the many actions of BDNF and to be a dorsal horn neuronal population participating in the sensory gating effects of ES^54^.

The goal of this study was to reveal mechanistic actions of strategies that improve locomotor outcomes after SCI at the level of identified spinal interneurons. We compared the individual and combined effects of AAV-BDNF and daily sub-motor threshold ES on locomotor recovery in an adult mouse model of complete SCI. We found that AAV-BDNF resulted in significant and sustained improvements in hindlimb stepping, but also exacerbated hyperreflexia compared to untreated SCI counterparts, replicating prior reports in rat^23,24,26^. BDNF-induced hyperreflexia was reduced during concurrent ES, but daily stimulation had no lasting effect on locomotor outcomes in mice with or without BDNF. Due to their role in gating proprioceptive afferents during locomotion, cellular properties of inhibitory RORβ neurons were explored as a possible point of control over locomotor circuitry. Using hierarchical clustering, we found that measures of cellular excitability of RORβ neurons correlated with the final functional outcome. Chemogenetic excitation of inhibitory RORβ neurons below the lesion also reduced hyperreflexia in mice overexpressing BDNF, mimicking the effects of ES *in vivo.* Taken together, our findings identify inhibitory RORβ neurons as an important target population to engage and modulate during the restoration locomotor function after complete SCI.

## Results

### AAV5-BDNF enables functional improvements in hindlimb stepping after complete SCI

We assessed and compared locomotor performance in adult RORβcre;tdTomato mice that were either untreated (SCI) or received intraspinal AAV-BDNF (SCI+BDNF) after a complete thoracic spinal transection at T8/9. Hindlimb step events on the treadmill in SCI+BDNF mice were significantly increased compared to untreated SCI counterparts (Figure 1A) from the first week post-SCI (Mann-Whitney test, p<0.001) and maintained throughout the time course. Improvements in hindlimb stepping were also evident in overground locomotion, assessed by the nine-point Basso Mouse Scale (BMS)^55^, where we found a significant increase in BMS scores of SCI+BDNF mice compared to SCI mice at all timepoints tested (Figure 1B, Mann-Whitney tests, p<0.001), with a time course similar to that seen with total step events. Whereas hindlimb movements in untreated SCI mice were restricted to ankle flexions with little to no engagement of knee or hip joints (Figure 1C) plantar steps comprised the majority of all treadmill hindlimb movements at later timepoints in SCI+BDNF mice (Figure 1D). These data demonstrate that below injury level AAV-BDNF is sufficient to elicit sustained, alternating, weight-supported plantar stepping after complete transection in adult mouse.

**Figure 1.**
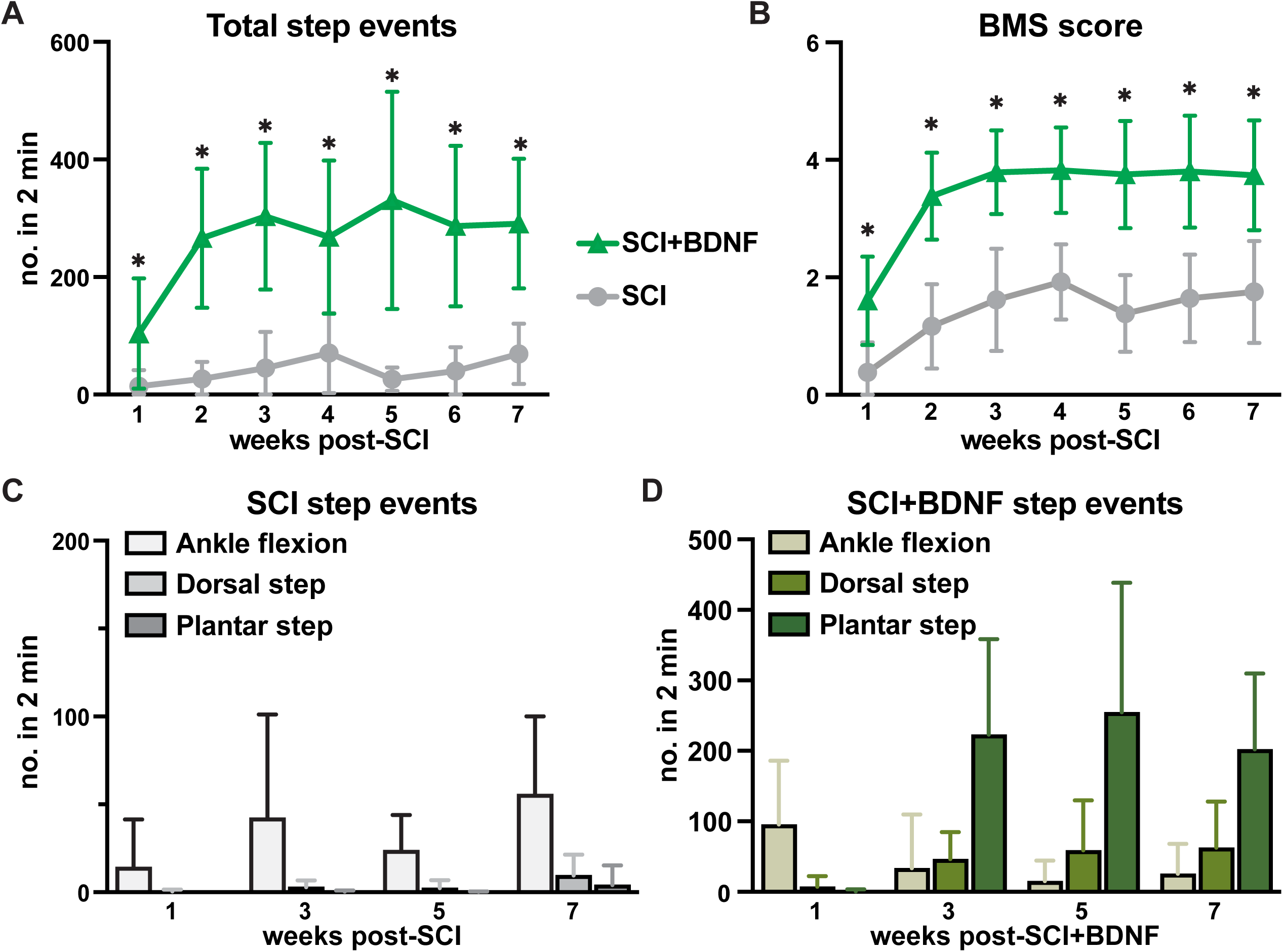
AAV5-BDNF enables functional improvements in hindlimb stepping after complete SCI. (A) Quantification of step-like movements, which include the number of closures of ankle joints, dorsal steps, and plantar steps observed while suspended over a moving treadmill during a 120s time window, and compared between SCI (gray) and SCI+BDNF groups (green, unpaired t-tests or Mann-Whitney tests, p<0.01). (B) Overground locomotor assessment in the open field using the nine-point Basso Mouse Scale (BMS, unpaired t-tests or Mann-Whitney tests, p<0.01). (C-D) Qualitative break down of total step-like movements performed by (C) SCI mice and (D) SCI+BDNF mice. Note the expanded y-axis in D. N=13 for SCI mice at all weeks except week 7 (N=11). N=20 for SCI+BDNF mice at all weeks except at week 3 (N=19), week 4 (N=17), week 6 (N=19) and week 7 (N=19). Data is represented as mean ± SD.

### Hyperreflexia is concomitant with BDNF-induced locomotor improvements

Previous SCI studies have explored the effects of AAV-BDNF in larger animal models and found increased measures of pain and/or spasticity alongside locomotor improvements^22-25^. Therefore, we sought to characterize aberrant hindlimb activity in SCI mice that received AAV-BDNF compared to untreated counterparts. In the early period of recovery after transection and injection of AAV-BDNF (weeks 1-2), as mice regained hindlimb weight-support, bouts of scoliosis, toe flaring/clasping, clonus, and hyperflexion in one or both hindlimbs emerged during treadmill and overground stepping (Figure 2A). The amount of time that SCI+BDNF mice spent exhibiting one or more signs of hyperreflexia during weekly treadmill sessions is reflected in the hyperreflexia score, which was significantly increased compared to SCI counterparts from the 2^nd^ week after injury (Figure 2B, Mann-Whitney tests, p<0.001) and until the last measurement 7 weeks post-SCI (unpaired *t*-test, p<0.001). Despite significantly higher hyperreflexia which often precluded stepping in at least one hindlimb, we found that step events, plantar steps, and BMS scores were not reduced with time (Figure 1). These findings indicate that, in addition to increased functional hindlimb stepping, AAV-BDNF increased hyperreflexia during treadmill locomotion.

**Figure 2.**
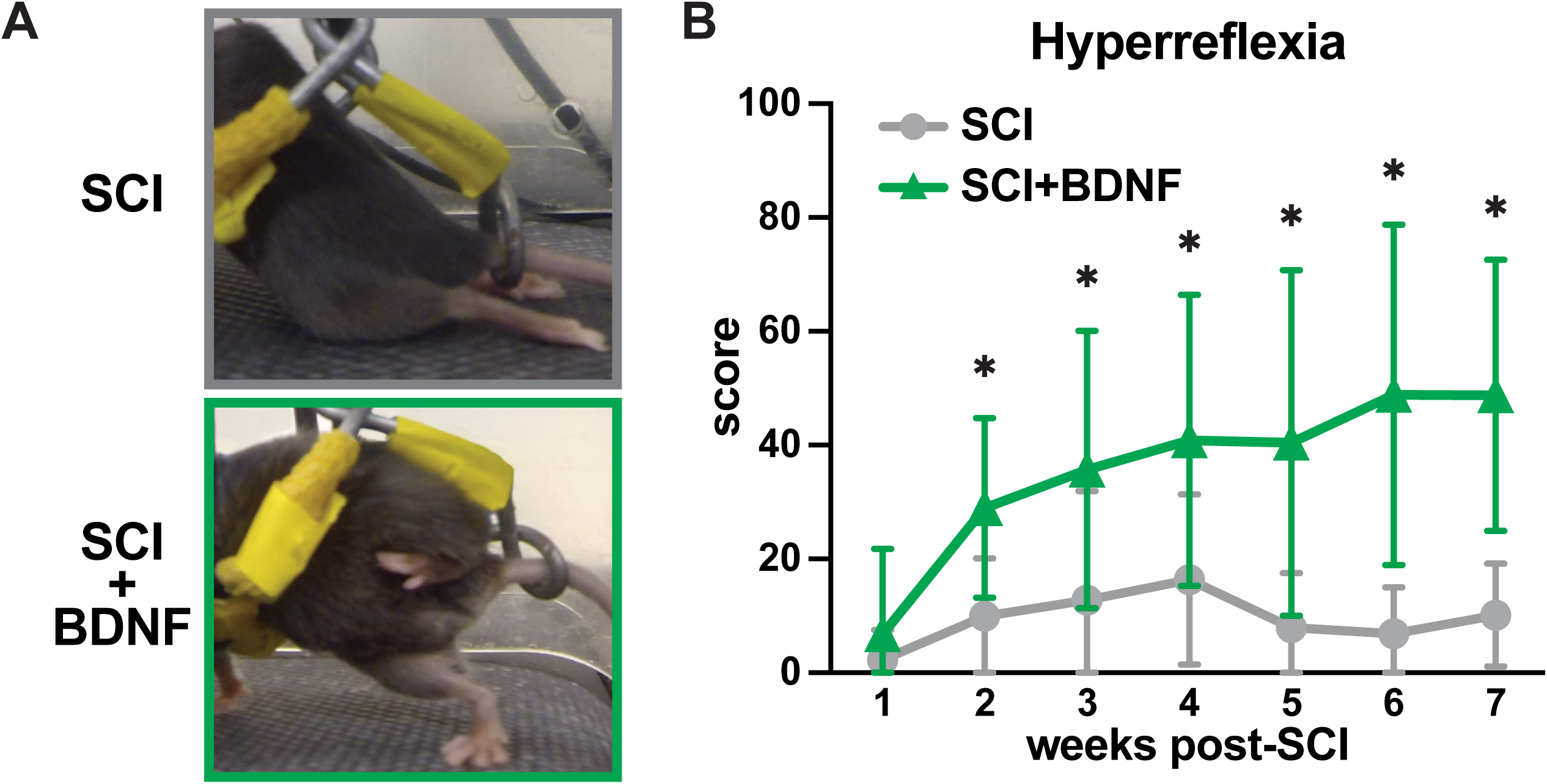
Hyperreflexia is concomitant with BDNF-induced locomotor improvements. (A) Video stills of an SCI+BDNF mouse captured during the 120s treadmill scoring period displaying dragging of hindlimbs one week after transection (gray outline), as typically seen in SCI mice, compared to the BDNF-induced stepping with the appearance of hyperflexion at later timepoints (green outline). (B) Quantification and comparison of weekly hyperreflexia scores between SCI (gray line) and SCI+BDNF (green line) mice (unpaired t-tests or Mann-Whitney tests, p<0.01). N values are the same as in Figure 1. Data is represented as mean ± SD.

### Daily ES mediates an acute improvement in BDNF-induced treadmill locomotion that is not sustained

Sub-motor threshold ES has been associated with increases in functional locomotion as well as decreases in hyperreflexive symptoms after SCI^5,56^ In order to determine whether ES alone had an effect on locomotor outcomes post-SCI in our model, we assessed treadmill locomotor performance in an additional set of SCI mice that received daily ES alone (SCI+ES). We first compared weekly measures of treadmill locomotion taken before ES sessions (SCI+ES^OFF^) to those taken during stimulation (SCI+ES^ON^) for individual mice using two-way repeated measures ANOVA. We found no differences in total step events (F^(1, 21)^=2.01, p=0.17, Figure 3A) nor hyperreflexia scores between conditions (F^(1, 21)^=1.58, p=0.22, Figure 3B). To determine whether daily ES treatment improved treadmill performance over time, we compared final measures taken from SCI mice against SCI+ES^OFF^ values and found no difference in total step events (Figure 3C, unpaired t-test, p=0.30) nor hyperreflexia scores (Figure 3D, unpaired t-test, p=0.29). Although we expected to see modest improvements in hindlimb function compared to untreated SCI counterparts, these data indicate a lack of effect from sub-motor threshold ES during stimulation and following prolonged ES training (>6 weeks).

**Figure 3.**
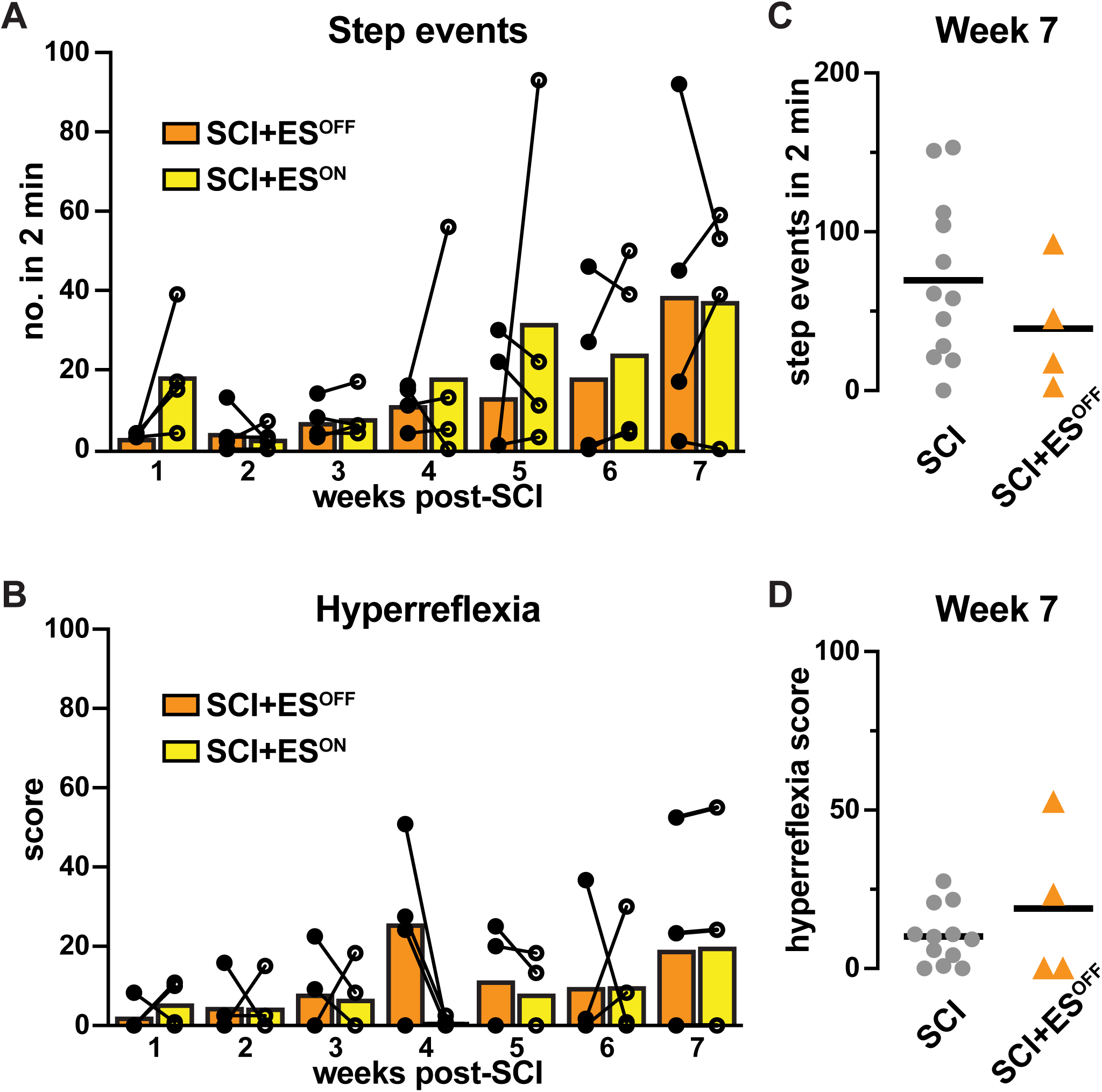
Sub-motor threshold ES has no effect on locomotor performance after complete SCI in mouse. Within-subjects comparisons of (A) total step attempts and (B) hyperreflexia scores in SCI+ES mice quantified with ES off (SCI+ES^OFF^, orange, closed circle) and ES on (SCI+ES^ON^, yellow, open circles; two-way RM ANOVA, Fisher’s LSD, p=0.17 and p=0.22, respectively). Comparison of locomotor outcomes between SCI and SCI+ES mice that received >6 weeks of daily ES using final measures of (C) total step-like movements (unpaired t-test, p=0.30 and (D) hyperreflexia scores (unpaired t-test, p=0.29).

Given its capacity to modulate sensorimotor signaling, we hypothesized that sub-motor-threshold ES would reduce BDNF-induced hyperreflexia and maximize functional recovery after SCI. Therefore, a separate group of SCI mice received AAV-BDNF in addition to daily sub-motor threshold ES at lumbar (L2) spinal segments while hindlimbs were suspended over the treadmill (SCI+BDNF+ES). We first compared measures of treadmill locomotion from videos taken before daily ES (SCI+BDNF+ES^OFF^) against those taken during ES the next day (SCI+BDNF+ES^ON^) to determine whether sub-motor threshold ES mediated a beneficial effect in BDNF-treated mice.

A two-way repeated measures ANOVA of hyperreflexia scores in individual mice under both conditions (SCI+BDNF+ES^OFF^ vs. SCI+BDNF+ES^ON^) found significant effects in condition (F^(1, 95)^=42.32, p<0.001), time (F^(6, 95)^=9.07, p<0.001), and interaction (F^(6, 95)^=2.43, p=0.031). Post hoc comparisons (unprotected Fisher’s LSD) revealed a significant reduction in hyperreflexia scores during sub-motor threshold ES at weeks 2 (p=0.03), 3 (p=0.002), 4 (p<0.001), 5 (p<0.001), and 7 (p=0.006) post-SCI (Figure 4A).

**Figure 4.**
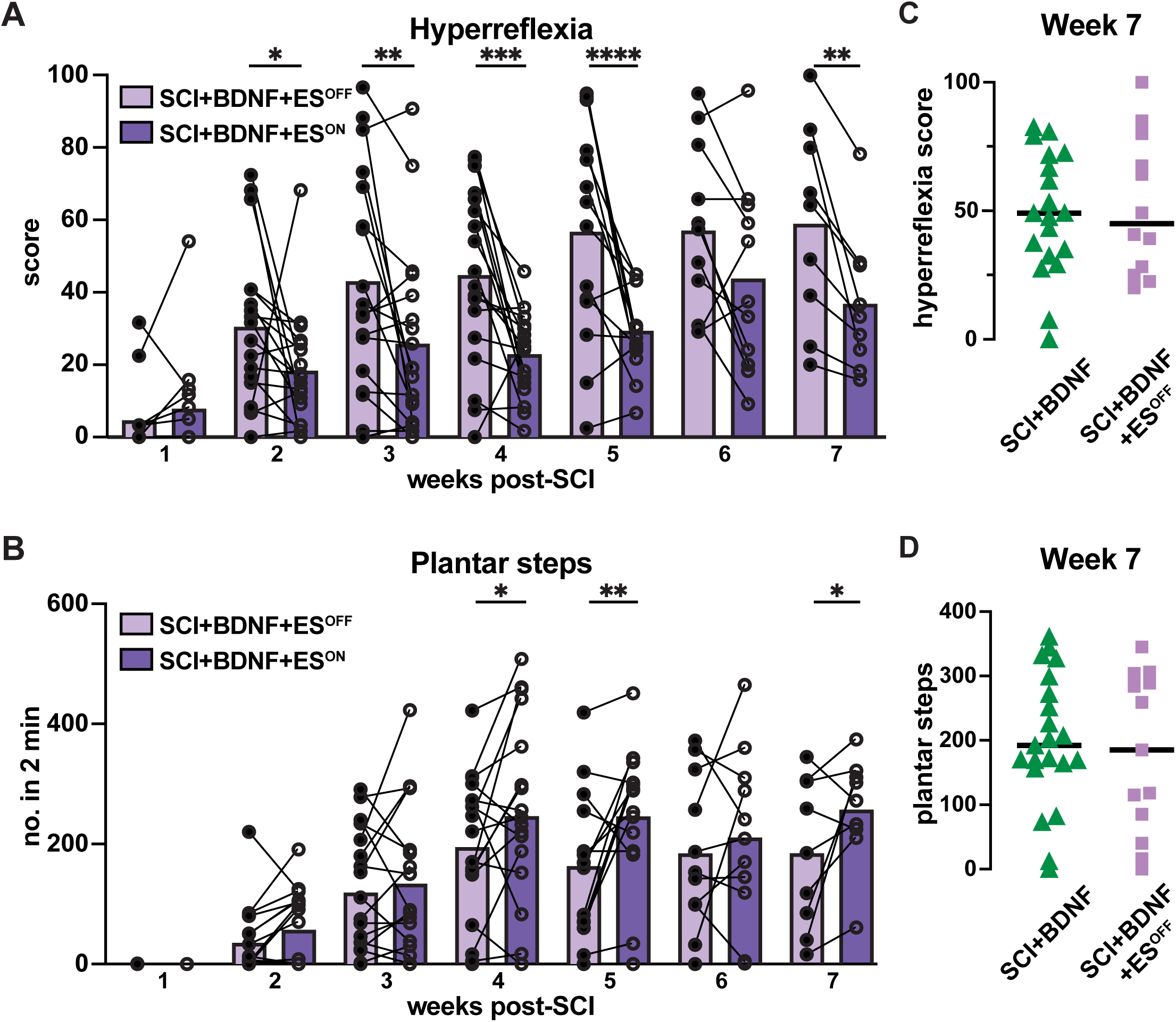
Daily ES mediates an acute improvement in BDNF-induced treadmill locomotion that is not sustained. Within-subjects comparisons (two-way RM ANOVA, Fisher’s LSD) of (A) functional plantar stepping and (B) hyperreflexia scores in SCI+BDNF+ES mice with stimulation off (SCI+BDNF+ES^OFF^, periwinkle, closed circles) and stimulation on (SCI+BDNF+ES^ON^, violet, open circles). (C-D) Comparisons of locomotor outcomes between SCI+BDNF and SCI+BDNF+ES mice that received >6 weeks of daily ES using final measures of (C) hyperreflexia scores (unpaired t-test, p=0.75) and (D) plantar steps (unpaired t-test, p=0.87). *p<0.05, **p<0.01, ***p<0.001, ****p<0.0001.

We performed similar comparisons for plantar stepping and found significant effects of condition (F^(1, 95)^=16.14, p<0.001) and time (F^(1, 95)^=12.86, p<0.001) but not in the interaction (F^(6, 95)^=1.51, p=0.184). In tandem with the reduction in hyperreflexia scores, we found significant increases in the quantity of plantar steps during stimulation at weeks 4, 5 and 7 post-SCI (Figure 4B, unprotected Fisher’s LSD, p=0.02, 0.001, and 0.02, respectively). Taken together, these findings indicate that sub-motor threshold ES alone has no effect on locomotor outcomes, but when combined with AAV-BDNF, there is an acute ES-mediated enhancement in locomotor function, likely through a decrease in hyperreflexia.

When combined with an additional source of excitatory modulation, ES has been shown to augment sensorimotor signaling and facilitate locomotor improvements during rehabilitation^57,58^. We hypothesized that, over time, ES would result in lasting improvements that would enhance functional outcomes in the absence of stimulation, beyond those achieved by BDNF alone. Therefore, we compared terminal measures of treadmill locomotion from SCI+BDNF mice against those from SCI+BDNF+ES^OFF^ mice to determine sustained differences in functional outcomes following combined treatment. However, we found no differences between BDNF-treated groups when comparing final measures of hyperreflexia scores (Figure 4C, unpaired t-test, p=0.75) and plantar steps (Figure 4D, unpaired t-test, p=0.87). Thus, as with SCI+ES mice, sub-motor threshold ES sessions had no lasting or cumulative effects on locomotor outcomes in mice overexpressing BDNF.

### Experimental mice cluster according to terminal functional outcomes

Our overarching goal was to investigate cell-specific changes to spinal locomotor circuitry that were associated with functional recovery and the therapeutic treatments tested. Contrary to our initial hypothesis, we found no lasting differences in functional outcomes between SCI+BDNF and SCI+BDNF+ES groups. Importantly, slice electrophysiology recordings occurred the day after the final ES session, reflecting the SCI+BDNF+ES^OFF^ condition. Additionally, we noted variability within groups for both stepping and hyperreflexia (see Figure 4C and 4D). Rather than comparing measures of cellular excitability between our initial behavioral treatment groups, we implemented unbiased clustering methods to identify functional groupings derived from locomotor and hyperreflexia scores measured at terminal timepoints irrespective of post-SCI treatment paradigm.

We considered the final (week 7) measures of ankle flexions, dorsal steps, plantar steps, and hyperreflexia as a set of parameters to identify functionally distinct clusters amongst all experimental mice which had associated patch clamp data (Figure 5A). We first performed a correlation analysis on these parameters to examine highly correlated variables^59^. There was an inverse correlation between ankle flexions and plantar steps; however, both variables remained in the analysis to account for the degree of recovery attained across all experimental mice. Agglomerative clustering approaches established 3 clusters which were evaluated by silhouette analyses (Figure 5B) and visualized in the construction of a dendrogram (Figure 5C).

**Figure 5.**
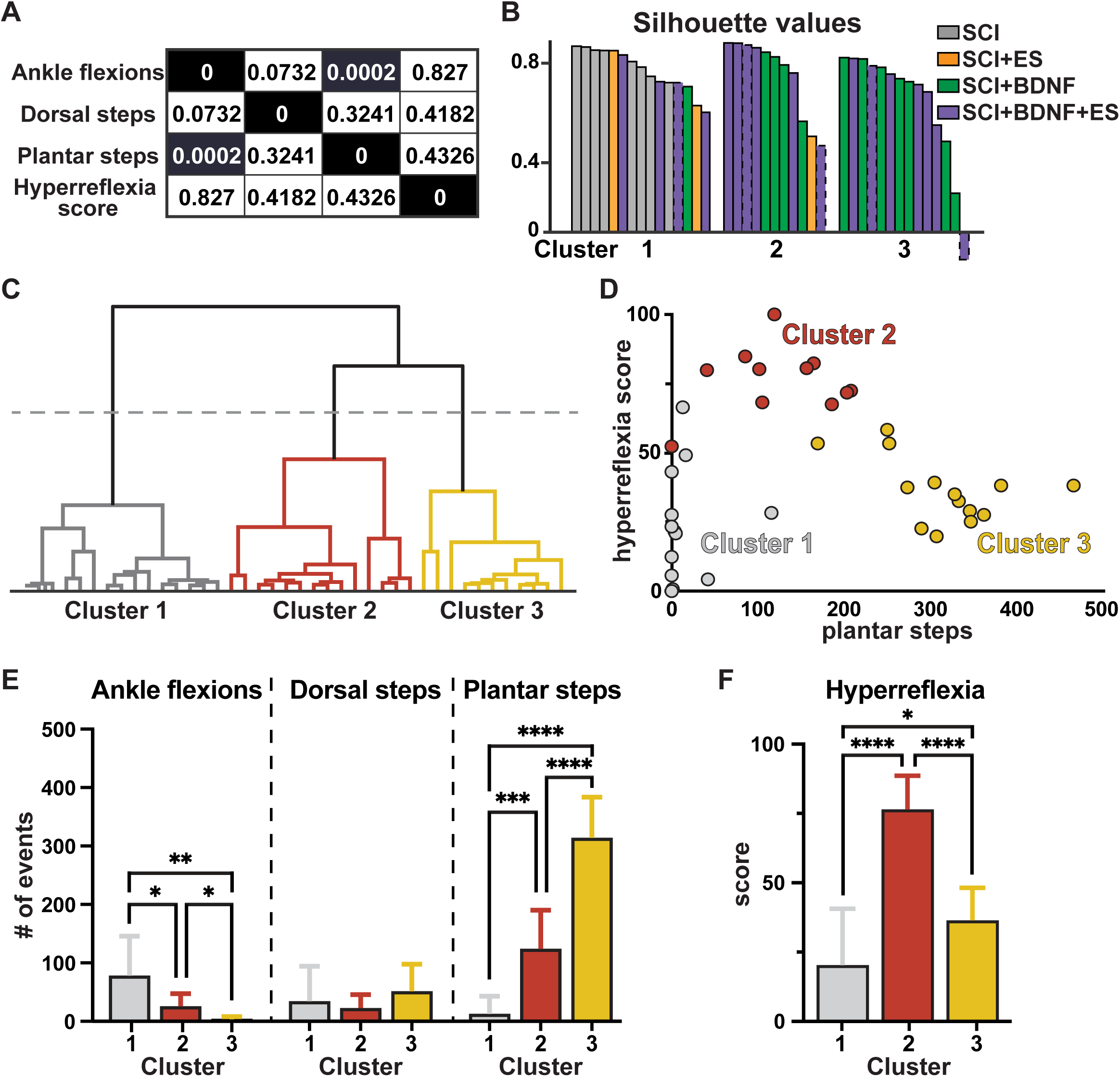
Experimental mice cluster according to functional outcomes. (A) Correlation coefficients for the 4 behavioral measures at terminal timepoints that were considered for cluster analyses. (B) Silhouette values for each mouse plotted in clusters, with each bar colored according to initial treatment paradigms received (gray=SCI, orange=SCI+ES, green=SCI+BDNF, purple=SCI+BDNF+ES). Red borders denote SCI+BDNF+ES mice which received >6 weeks of ES post-SCI. (C) Dendrogram plot identifying 3 clusters by natural divisions (colored throughout as gray, red, and gold). (D) Scatter plot displaying an inverted U when comparing plantar steps (X-axis) against hyperreflexia score (Y-axis). (E) Comparison of individual step measures to display functional differences between clusters. (F) Terminal hyperreflexia scores compared between clusters (one-way ANOVA, Tukey’s *post hoc* test, p<0.01). *p<0.05, **p<0.01, ***p<0.001, ****p<0.0001. Data is represented as mean ± SD.

Cluster compositions showed that Cluster 1 was mixed as far as treatment groups (visualized Figure 5B). Clusters 2 and 3 contained primarily mice which received BDNF (SCI+BDNF and SCI+BDNF+ES), with only one SCI+ES mouse included in Cluster 2. Clustered data points were generally separable and distributed along an inverted-U on a plot of plantar steps vs. hyperreflexia score (Figure 5D). Cluster 1 contained all untreated SCI mice (N=8) along with mice from every other SCI+treatment group, corresponding to the gray points in Figure 4D. These mice typically had very low plantar stepping. Cluster 2 was comprised of a single SCI+ES mouse (N=1), among SCI+BDNF (N=4) and SCI+BDNF+ES mice (N=4), corresponding to the red points in Figure 5D. These mice had high hyperreflexia but were capable of some plantar stepping. Cluster 3 consisted only of mice that received AAV-BDNF, whether it be alone (N=7) or in combination with daily ES (N=5), corresponding to gold points in Figure 5D. These mice had higher plantar stepping and lower hyperreflexia (see below) and were mostly the highest performing mice. Five mice that received ES until the wires failed after quantifications in week 3 (N=3) or 5 (N=2) were included in this analysis as well, one of which was in Cluster 1 whereas the others were equally distributed in Cluster 2 (3 weeks ES, N=1, 5 weeks ES, N=1) and Cluster 3 (3 weeks ES, N=1, 5 weeks ES, N=1).

We compared the clusters statistically on the individual treadmill step measures used as input into the cluster analysis to determine functional differences between clusters (Figure 5E-F). We found significant differences in ankle flexions (Welch’s one-way ANOVA, W^(2, 16)^= 13.85, p<0.001) and plantar steps (Welch’s one-way ANOVA, W^(2, 19)^= 111.6, p<0.001). Mice in Cluster 1 rarely performed plantar steps but had a significantly higher number of ankle flexions compared to both Cluster 2 (Tamhane’s post hoc, p=0.03) and Cluster 3 (Tamhane’s post hoc, p=0.002). Mice in Cluster 2 performed significantly more plantar steps than mice in Cluster 1 (Tamhane’s post hoc, p<0.001), but significantly fewer plantar steps than mice in Cluster 3 (Tamhane’s post hoc, p<0.001). A comparison of hyperreflexia scores indicated significant differences between clusters (Figure 5F, one-way ANOVA, F^(2, 31)^= 41.97). Mice in Cluster 3 had a significantly higher hyperreflexia scores than mice in Cluster 1 (36% vs. 20%, p=0.02), but mice in Cluster 2 had a significantly higher hyperreflexia score (77%) than mice in both other clusters (compared to Cluster 1, p<0.001; compared to Cluster 3, p<0.001). Thus, Clusters 1, 2, and 3 can be considered to correspond to the following descriptions: no stepping//low hyperreflexia, moderate stepping//high hyperreflexia, and high stepping//moderate hyperreflexia, respectively. This data demonstrates the possibility for experimental mice to be clustered based on motor phenotype and shows that motor phenotype does not directly correspond to post-SCI treatment paradigm in that Cluster 2 and 3 both contained mice that received either SCI+BDNF and SCI+BDNF+ES treatments and Cluster 2 had 1 SCI+ES mouse.

### Electrophysiological properties of inhibitory RORβ neurons correlate with post-SCI functional locomotor outcomes

Whereas non-specific excitatory modulation is sufficient to reactivate silent motor pathways, spinal inhibitory mechanisms can facilitate gating to bias relevant sensorimotor signaling to promote functional improvements after SCI. We thus hypothesized that inhibitory mechanisms were likely better engaged in Cluster 3 mice. Although AAV-BDNF and ES are likely to have numerous actions on the spinal sensorimotor system, we focused on interneurons involved in gating proprioceptive sensory pathways which could be recruited to enhance locomotor function by reducing hyperreflexia. As shown in Figures 1 and 2, AAV-BDNF leads to a more excitable locomotor circuitry (stepping) but also increases the excitability sensorimotor pathways leading to varying degrees of hyperreflexia in individual mice. Excitation, particularly when excessive or aberrant, could be balanced by spinal mechanisms of inhibition, if appropriately engaged. Interestingly, the features of hyperreflexia observed in our BDNF mice resembled that of the ‘duck gait’ phenotype characterized in mutant RORβ mice ^51^. Given that RORβ neurons express TrkB receptors and mediate presynaptic inhibition of proprioceptive afferents during locomotion ^51^, we thus focused on deep dorsal inhibitory RORβ neurons to determine their cellular excitability in relation to post-SCI locomotor clusters and outcomes. Membrane and firing properties were recorded from tdTomato-expressing RORβ neurons in the medial region of the deep dorsal horn and analyzed prior to our functional clustering analysis.

Consistent with prior studies of inhibitory populations^60,61^, we found that most inhibitory RORβ neurons from uninjured cords displayed tonic firing patterns (89%) in response to relatively low levels of injected current (mean rheobase = 10.6 +/-5.4). The other RORβ neurons displayed either initial burst or delay firing, both occurring with low incidence (Figure 6A). The proportion of neurons from Cluster 1 firing tonically during a current step (73%) was significantly lower than the proportion of tonically firing neurons from Cluster 2 (94%, Fisher exact test, p=0.015) and Cluster 3 (100%, Fisher exact test, p=0.001). Although the proportion of tonic firing patterns observed in Cluster 1 was not significantly different compared to uninjured neurons (Fisher exact test, p=0.07), we found that single spike neurons were exclusive to Cluster 1, the cluster comprised of mice with the lowest hindlimb activity (Figure 6A, B). Additionally, during continuous recordings at resting membrane potential, a higher proportion of neurons in the best stepping mice, Cluster 3 (93%), was spontaneously firing action potentials compared to those from uninjured mice (71%, Fisher exact test, p=0.028) and Cluster 1 (71%, Fisher exact test, p<0.024, Figure 6C). We found no significant differences in the frequency of spontaneous firing between any group when it occurred (Figure 6D). These recordings show that adult deep dorsal RORβ neurons mainly display tonic firing patterns. When considering functional cluster designations, neurons from the best stepping SCI mice (Cluster 3) exhibited increased numbers of neurons with spontaneous spiking activity compared to those from the clusters of relatively more disabled SCI counterparts.

**Figure 6.**
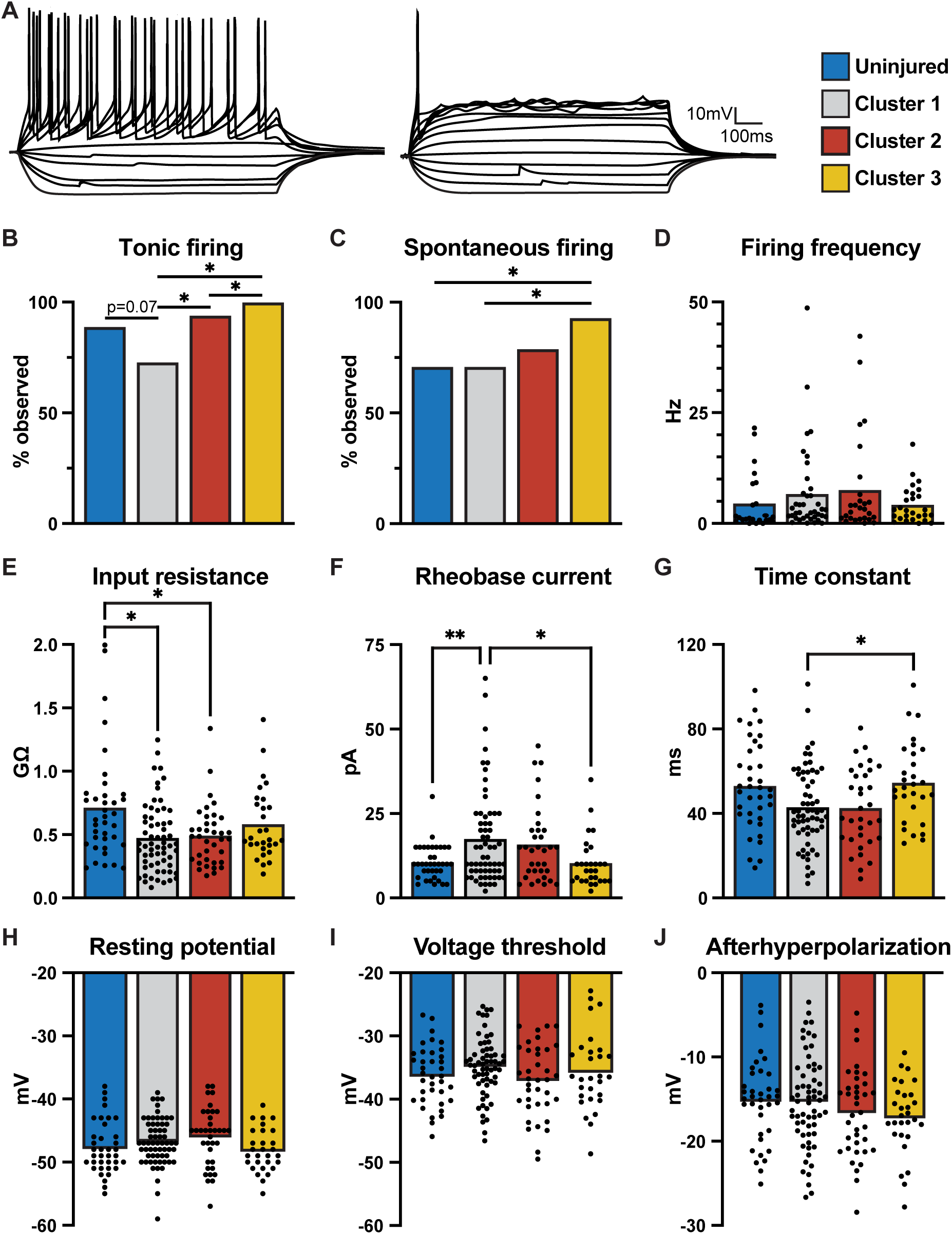
Electrophysiological properties of inhibitory RORβ neurons correlate with post-SCI functional outcomes. (A) Examples of tonic (left) and non-tonic (right) firing patterns of adult RORβ neurons in response to injected current steps. Neurons that fired with an initial burst or delay, but transiently, were categorized with single spike neurons as non-tonic. (B) The proportion of evoked firing patterns observed between cells from uninjured (blue) mice or SCI mice in Cluster 1 (gray), Cluster 2, (red), or Cluster 3 (yellow; Fisher exact test, p<0.01). Colors remain constant throughout. (C) Proportion of neurons that fired spontaneously at resting membrane potential (Fisher exact test, p<0.01). (D-F) Comparisons of intrinsic RORβ neuron properties (one-way ANOVA with Tukey’s post hoc or Welch’s with Tamhane’s T2 post hoc). We found significant differences in (D) membrane input resistance (Welch’s ANOVA, p=0.017), (E) rheobase current (Welch’s ANOVA, p=0.001), and (F) time constant (one-way ANOVA, p=0.008) when comparing group mean values. Post hoc tests are described in Methods and listed in the text, *p<0.05, **p<0.01.

We next compared other active and passive measures of cellular excitability to determine if there were differences in the intrinsic properties of RORβ neurons from uninjured and functional clusters (Figures 6E-J). Group mean comparisons identified significant differences in input resistance (Figure 6E, Welch’s ANOVA, W^(3, 76)^= 3.6, p=0.017), rheobase (Figure 6F, Welch’s ANOVA, W^(3, 77)^= 35.8, p=0.001), and time constant (Figure 6G, one-way ANOVA, F^(3, 156)^= 3.6, p=0.043) which were followed by post hoc comparisons. Compared to RORβ neurons from uninjured cords, input resistance was significantly lower in neurons from lower stepping Clusters 1 and 2 (Tamhane’s post hoc, p=0.02 and p=0.04, respectively) but not in neurons from the cords of the best stepping mice in Cluster 3 (Tamhane’s post hoc, p=0.60). Rheobase current was significantly higher in cells from mice in low hindlimb activity Cluster 1 compared to those from uninjured and Cluster 3 (Tamhane’s post hoc, p=0.004 and p=0.01, respectively). Additionally, membrane time constant in RORβ neurons was significantly longer in cells from mice in best stepping Cluster 3 compared to those in low hindlimb activity Cluster 1 (Tukey’s post hoc, p=0.04). Despite the shift in proportion of neurons firing spontaneously at rest described above, we found no difference in resting membrane potentials between all groups (Figure 6H, F^(3, 160)^=1.85, p=0.14). No differences were found when comparing action potential voltage threshold (Figure 6I, F^(3, 160)^=1.53, p=0.21), nor afterhyperpolarization (Figure 6J, F^(3, 160)^=1.38, p=0.25). The properties of RORβ neurons were similar in the groups with the best stepping, uninjured and Cluster 3. These findings provide further evidence to suggest a correlation between post-SCI locomotor function and increased measures of cellular excitability in deep dorsal RORβ neurons.

### Chemogenetic excitation of RORβ neurons improves BDNF-induced treadmill locomotion *in vivo*

Given the correlations between the cellular excitability of deep dorsal RORβ neurons and functional outcomes after SCI and that ES decreases hyperexcitability during stimulation, we sought to determine whether activation of RORβ neurons could mimic the beneficial effects observed during ES in SCI+BDNF mice. We used a cre-dependent Gq DREADD-based strategy to specifically activate RORβ neurons in SCI+BDNF mice (SCI+BDNF+RORβ^DREADD^). A separate group of RORβcre mice was injected with a reporter construct (SCI+BDNF+RORβ^mCherry^) and served as controls. Histological preparations in a subset of experimental mice revealed mCherry expressing neurons immediately below the transection and in spinal segments 4-6mm caudally (Figure 7A). We quantified the mean number of neurons that expressed mCherry, and the percentage of these that also expressed inhibitory neuron marker Pax2 in the medial deep dorsal horn (mDDh), the superficial dorsal horn (SDh), and other regions (Other) across at least 10 sections (Figure 7B). The highest concentrations of mCherry^+^ neurons were located in the SDh and mDDh, regions where RORβ neurons are expected^51^. Also consistent with prior studies^51,62,63^, the majority of mDDh neurons and approximately half of the SDh neurons expressed Pax2, indicating that they are inhibitory^64^.

**Figure 7.**
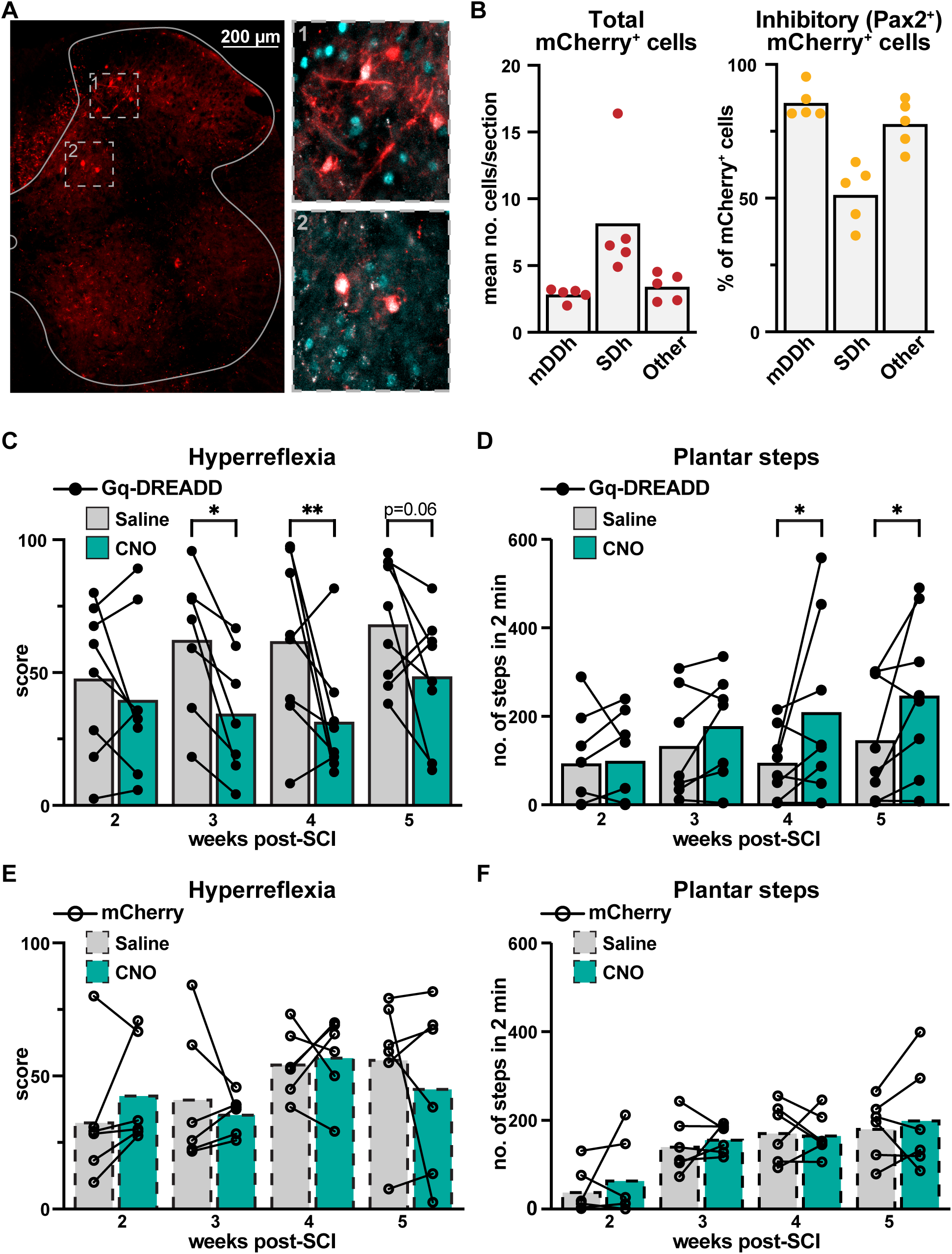
Chemogenetic excitation of RORβ neurons improves BDNF-induced locomotion *in vivo.* (A) A transverse section of the lumbar spinal cord from an experimental SCI+BDNF+RORβ^mCherry^ mouse 5 weeks after transection with mCherry^+^ cells in red and Pax2^+^ cells in cyan. Insets (1, 2) are denoted with dotted lines. (B) Quantification of mCherry^+^ and mCherry^+^/Pax2^+^ cells in the medial deep dorsal horn (mDDh), superficial dorsal horn (SDh), or other regions of the cord in SCI+BDNF+RORβ^DREADD^ mice. Weekly comparisons of (C) hyperreflexia scores and (D) plantar step for individual SCI+BDNF+RORβ^DREADD^ mice following injection of vehicle (gray) or CNO (teal). The same comparisons of hyperreflexia scores (E) and plantar steps (F) for individual SCI+BDNF+RORβ^mCherry^ mice following injection of vehicle (gray dashed) or CNO (teal dashed). The same individual mice were quantified each week in both conditions (vehicle and CNO), with the display highlighting the comparisons within a given week, overlayed on group mean values (two-way RM ANOVA, Fisher’s LSD, p<0.05). Post hoc tests as described in Methods and listed in the text, *p<0.05, **p<0.01.

We compared weekly behavioral assessments after administering either vehicle (saline) or CNO (5mg/kg) on separate, but sequential, testing days to determine the effects of RORβ chemogenetic activation on hyperreflexia and locomotor measures. In SCI+BDNF+RORβ^DREADD^ mice, we found a significant difference between conditions when comparing plantar step quantifications (two-way RM ANOVA, p=0.005). After CNO, SCI+BDNF+RORβ^DREADD^ mice performed significantly more plantar steps at post-SCI weeks 4 and 5 compared to vehicle (Fisher’s LSD, p=0.014 and p=0.028, respectively). Hyperreflexia scores were also significantly different between conditions (two-way RM ANOVA, F^(1, 27)^=17.30, p<0.001). Post hoc comparisons revealed a significant reduction in hyperreflexia scores at post-SCI weeks 3 and 4 in the presence of CNO compared to saline (Fisher’s LSD, p=0.016 and p=0.006, respectively). In SCI+BDNF+RORβ^mCherry^ mice, we found no differences in plantar stepping (F^(1, 20)^= 0.43, p=0.52) nor hyperreflexia scores (F^(1, 20)^= 0.36, p=0.85) between CNO and vehicle conditions, demonstrating no behavioral differences in response to CNO. The results from this experiment demonstrate that, similarly to ES, chemogenetic activation of RORβ interneurons alone reduces hyperreflexia *in vivo* and improves treadmill locomotion in SCI+BDNF mice.

## Discussion

Our goal was to identify cell-specific spinal plasticity occurring following chronic SCI and resulting from locomotor improvement strategies in order to determine mechanisms which can be further leveraged to promote functional recovery and prevent maladaptive changes. Molecular profiling has expanded the number of distinct types of spinal interneurons^65,66^, although few of these interneuron populations have been examined in the context of SCI or ascribed a role in the process of locomotor recovery after SCI^67-72^. We used virally-delivered BDNF to promote sustained stepping in mice after complete SCI; however, as reported elsewhere, there was a concomitant increase in hyperreflexia^23,24,26^. Sub-motor-threshold epidural stimulation decreased hyperreflexia and improved stepping but only during ES delivery, suggesting an acute inhibitory effect of ES which dampens the hyperexcitable reflexes. We found that outcomes of treatment overlapped between BDNF alone and BDNF+ES and formed two clusters with differing levels of hyperreflexia as one parameter driving clustering. Although there are undoubtedly several populations of spinal interneurons involved, we chose to first focus on RORβ interneurons in the deep dorsal horn because they are involved in proprioceptive gating, they express TrkB receptors, and removal of TrkB receptors from RORβ neurons results in hyperflexion during walking^51^. Measures of cellular excitability in inhibitory RORβ interneurons correlated with functional outcome clusters in SCI mice. The SCI cluster with the best locomotor outcomes, composed exclusively of SCI+BDNF and SCI+BDNF+ES mice, also exhibited the RORβ neurons with the highest excitability (lowest rheobase) and highest proportion spontaneously active compared to other SCI groups, and intrinsic properties that were indistinguishable from uninjured mice. Further, BDNF-induced hyperreflexia was reduced and stepping was modestly improved during chemogenetic excitation of RORβ neurons, results very similar to those observed with ES treatment. Taken together, we identify deep dorsal inhibitory RORβ IN as a point of control to both reduce hyperreflexia and improve locomotor outcomes after complete SCI.

### AAV-BDNF and ES enhance locomotor function following SCI in mouse

Brain derived neurotrophic factor (BDNF), signaling through the tropomyosin receptor kinase B (TrkB) receptor, has been implicated in functional recovery following various rehabilitation strategies in both animal models of SCI^11,13,17,73-75^ and in humans with SCI^10,76^. However, exogenous BDNF enhances afferent-evoked inputs in the dorsal and ventral horn and can facilitate a pro-nociceptive spinal state^77-79^. Virally overexpressing BDNF in spinal rats improves locomotor function but increases aberrant hindlimb activity^23,24,26^. Thus, BDNF has apparent treatment potential, but a greater mechanistic understanding will be necessary to facilitate functional recovery and mitigate detrimental effects. Our behavioral findings in mice align with these prior reports and our complete transect model also allows for determination of spinal mechanisms at play in isolation from descending influences.

The expression of the virally-delivered BDNF is continuous and non-specific, which is likely why both the promotion of stepping and hyperreflexia are observed. However, hyperreflexia may be beneficial in preventing muscle atrophy after SCI^80,81^, and some degree of spasticity can facilitate volitional movements^82-84^. This is consistent with the mouse behavioral data here (Figure 5) in which the best steppers (Cluster 3) exhibit moderate hyperreflexia. The lesser functioning groups (Clusters 1 and 2) have either very low or very high hyperreflexia. Thus, in the ideal case, BDNF expression may be targeted to particular regions or neuronal populations in a temporally controlled manner to facilitate locomotor improvements and provide the ‘right degree’ of hyperreflexia.

Unlike BDNF, ES is used clinically, yet little is known about the mechanisms of action^27,28,46^. Recent clinical findings demonstrate sustained improvements in standing or stepping performance after complete and chronic SCI that persists even in the absence of stimulation^42,85,86^. However, these results were obtained following significant motor rehabilitation. We did not observe enhanced chronic improvements with the addition of ES. The locomotor outcomes of SCI+BDNF and SCI+BDNF+ES groups were quite similar. This may be due to our stimulation paradigm. Unlike in larger animals such as rat^87^ or cat^88^, we were unable to evoke stepping with ES due to lower intensities of stimulation in order to avoid direct activation of motor neuron axons^89^ causing co-contraction of antagonistic muscles and occlusion of functional stepping. Sub-motor threshold ES facilitates spontaneous motor activity below complete spinal transection in adult rats during stimulation^90^, but we observed no difference in SCI mice lacking BDNF between conditions (SCI+ES^OFF^ vs. SCI+ES^ON^). Further optimization may be possible with a longer duration protocol or a multifrequency paradigm^72^, or pharmacotherapy in parallel^91^. In addition to direct effects on locomotor circuitry, low intensity ES has anti-spastic and anti-nociceptive effects^56,92^, likely via the recruitment of neuronal components involved in sensory gating^54,93,94^. This is consistent with our observed acute reductions in hyperreflexia during ES in SCI+BDNF+ES mice.

### Cell-type specific neuronal plasticity post-SCI and following intervention

In the context of SCI, a few populations of transmitter- or genetically-identified interneurons in the dorsal and ventral horns have been assessed. Intrinsic measures of cellular excitability were increased post-SCI in both lamina I inhibitory interneurons^95^ and in deep dorsal excitatory interneurons with bursting behavior in response to afferent stimulation^96^, with the latter population being associated with sensorimotor reflex responses^97^. In contrast, none of the studies examining the cellular and circuit properties of genetically-identified ventral interneuron populations have shown any changes in intrinsic properties post-SCI^68,69^. We found that deep dorsal inhibitory RORβ interneurons display decreases in measures of cellular excitability following chronic transection. Thus, in addition to the loss of descending drive and proprioceptive input to RORβ interneurons, a reduction in cellular excitability further compounds the disinhibition of spinal reflex pathways^8,98,99^.

### Deep dorsal inhibitory RORβ excitability correlates with functional outcomes *in vivo*

Given the variety of functions performed by the spinal cord and the diversity of interneurons involved^100-102^, combination treatments are likely to be critical to locomotor recovery following SCI. Previous studies using AAV-BDNF or ES have largely focused on motor neuron physiology and have indirectly characterized changes occurring at the interneuronal level through changes in neurotransmitter expression patterns^23-25,103^. However, current understanding of their effects at the level of genetically identified interneurons is limited, even more so regarding possible combinatorial effects.

Afferents are a primary target of both BDNF and ES, as with most post-SCI locomotor rehabilitation strategies^4,104^. Proprioceptive input is essential for functional locomotor recovery after injury^105,106^. Inhibitory RORβ neurons involved in sensory gating are activated upon afferent recruitment; therefore, RORβ activity should be linked with overall sensory feedback. Both BDNF and ES interventions are likely to act via multiple interneurons and pathways, but deep dorsal RORβ interneurons are well-positioned to mediate the effects of AAV-BDNF, through the expression of TrkB receptors, and ES, via their position in proprioceptive afferent pathways^51^. Accordingly, we were able to identify physiological changes to RORβ neurons which were linked with measures of treadmill performance in the experimental mice that the neurons were recorded from. Thus, amplifying input, via BDNF-related sprouting and synaptic strength changes^16^, and biasing towards activation of inhibitory control pathways with sub-motor threshold ES^107,108^ may be a way to preserve necessary activations while blocking aberrant input. Indeed, we were able to replicate the effects of ES by selectively activating RORβ neurons in the DREADD experiments.

Although sub-motor threshold ES did not have a lasting behavioral effect, it reduced hyperreflexia acutely during stimulation. Chemogenetic activation of RORβ neurons resulted in similar effects, both limiting hyperreflexia and improving stepping. This, together with the correlation of enhanced RORβ excitability and better stepping function, suggests that RORβ neurons are involved in locomotor functional recovery and can be acutely recruited to enhance sensory gating to restore inhibitory counterbalance in the absence of descending control. DREADD-mediated activation was not restricted to the deep dorsal population of RORβ neurons so we cannot rule out the possibility that the superficial RORβ neurons^51,62,109^ may also be involved. Thus, effects observed are not necessarily unique to this population. However, our results suggest that RORβ neurons are a population that can be targeted to limit hyperreflexia and assist in providing inhibitory counterbalance to therapeutic strategies that globally enhance neuronal excitability to restore locomotion after SCI.

Taken together, our results replicate that BDNF enhances locomotor circuits and sensory input, resulting in both locomotor improvements and hyperreflexia, and show that ES can modulate the hyperreflexia. In some treatments, the excitability of RORβ interneurons is enhanced (i.e., Cluster 3 mice) and this is associated with improved stepping and less hyperreflexia. In other cases, RORβ interneuron excitability is lower and the behavioral phenotype of the mice (Cluster 2) resembles that of mice with TrkB receptors removed from RORβ interneurons^51^. It is possible that the continuous overexpression of BDNF leads to a desensitization, saturation, and/or internalization of TrkB receptors^110-112^, any of which could explain the loss of beneficial effects in significant numbers of treated mice through increased hyperreflexia. Further, our results suggest the importance of temporal control and critical windows of opportunity when manipulating BDNF expression levels.

## Acknowledgments

We thank Dr. Jed Shumsky for consultations on statistics, Drexel ULAR staff members for expert animal care, and members of the Marion Murray Spinal Cord Research Center for discussions related to the work. We also thank Shayna Singh, Jenna McGrath and Chris West for comments on a version of the manuscript. This work was supported by NIH grants NS104194 (S.F.G. and K.J.D.), NS130799 (K.J.D.), and F31 NS127584 (N.J.S.), and the Drexel Dean’s Fellowship for Excellence in Collaborative or Themed Research (N.J.S.).

## Author Contributions

N.J.S., S.F.G., and K.J.D. conceptualized the study. N.J.S., S.J.A., and L.Y. performed behavioral experiments. N.J.S. performed the electrophysiology experiments. N.J.S. and J.H.W. performed the histology experiments. N.J.S., J.H.W., S.J.A., D.L.G-R., S.F.G., and K.J.D. analyzed the data. N.J.S. and K.J.D. wrote the manuscript with input from all authors.

## Methods

### Mouse lines and experimental groups

Experiments were performed using male and female RORβ::Cre; Rosa26-flox-stop-flox-tdTomato (Ai9) mice (from Jax mice, #023526 and #007909, respectively)^113,114^. All experimental procedures followed National Institutes of Health guidelines and were approved by the Institutional Animal Care and Use Committee at Drexel University. The experiments were performed in adult (>6 weeks) mice without any surgical procedures (uninjured mice *N*=19) and four different treatment conditions: (1) mice with complete spinal transection surgery and either no treatment (SCI mice, *N*=15), (2) mice with complete SCI and treatment with daily ES alone (SCI+ES mice, *N*=4), (3) complete SCI and treatment with AAV5-BDNF alone (BDNF mice, *N*=21) mice, or (4) complete SCI and treatment with daily ES and AAV5-BDNF in combination (SCI+BDNF+ES mice, *N=24*).

### SCI surgery

Mice (6-10 weeks old) were anesthetized with isoflurane (4% induction, 2% maintenance). Dorsal skin was shaved and sterilized with betadine and isopropyl alcohol. Incision was performed from the thoracic to lumbar vertebral segments. Following laminectomy, approximately one segment of spinal cord was removed, resulting in a complete transection of the thoracic spinal cord at T8-T10. Buprenorphine SR (0.5 mg/kg) and either ampicillin (20 mg/kg) or Baytril (10 mg/kg) were subcutaneously administered perisurgically. Mice were monitored and bladders were expressed manually twice daily. Completeness of transection and condition of the caudal spinal cord were visually assessed at dissection for electrophysiology or histology.

### Viral vector and intraspinal injection

AAVs were intraspinally injected to transduce neurons to overexpress our genes of interest (i.e., BDNF, hM3D) during the spinal transection surgery described above. The plasmid constructs containing the human BDNF encoding sequence and secretory signal sequence, under the control of a CMV promoter, were provided by Dr. Fred Gage (Salk Institute) and produced by the Penn Vector Core. BDNF mice received 0.25µL AAV5-BDNF (titer ∼2.5 x10^10^ viral particles/mL) injected bilaterally immediately following, and caudal to, the transection using a glass micropipette with Hamilton syringe (#80135) which was affixed to a micromanipulator and positioned in alignment with the caudal portion of the transected spinal cord. For chemogenetic manipulations *in vivo*, 0.25μL of AAV-DIO-hM3D(Gq DREADD)-mCherry or AAV-DIO-mCherry was co-injected with AAV-BDNF in separate subset of RORβcre mice. pAAV-hSyn-DIO-hM3D(Gq)-mCherry and pAAV-hSyn-DIO-mCherry were gifts from Bryan Roth ^115^ with viral preps purchased from Addgene (http://n2t.net/addgene:44361, RRID: Addgene 44361; http://n2t.net/addgene:50459; RRID:Addgene 50459). In all cases, the tip of the micropipette syringe was inserted approximately 200-250µm from the dorsal surface of the cord to a depth of 1-1.5mm and the syringe’s content was expelled over the course of 5 minutes and allowed an additional 5 minutes to diffuse into surrounding tissue. Peri-surgical treatment and monitoring were identical to SCI mice (see above).

### Wire placement for epidural stimulation

In experimental mice receiving ES, a partial laminectomy was performed above the spinal L1 segment during the same surgery and interlaminar incisions were made using fine scissors to allow for implantation of ES leads over the L2 segment. ES wires were constructed using two separate 0.002” Teflon insulated stainless steel wires (#793400, A-M systems, Sequim, WA) with offset exposures that were joined together with two-part epoxy at the implanted end (JB Kwik). An incision was made to expose the skull and allow the emerging leads to be threaded under the nape and soldered to a two-header pin connector headcap. Wires were sutured to the vertebral spine caudal to the insertion and muscle above the transection using nylon sutures before closing the incision. Two screws were implanted into the skull which the headcap was affixed to using dental cement. During ES, the headcap was connected to an optically isolated pulse stimulator (A-M Systems Model 2100, Sequim, WA) using two leads drawn from flexible ribbon cables.

### Locomotor rehabilitation and assessment

Starting one week prior to surgery, all experimental mice were placed on a moving treadmill (3 meters/minute) daily for 10 minutes to acclimate. Testing and/or ES began 1 week following injury and continued for 6 weeks. Mice lacking epidural leads (SCI and SCI+BDNF) remained in their home cage for this duration, except for weekly assessments. Mice treated with ES (SCI+ES and SCI+BDNF+ES) received sub-motor threshold stimulation (30µA, 200µs pulses at 40Hz) for 10 minutes a day, 5 days per week while suspended over a moving treadmill using a custom, elastic, supportive harness to position fore- and hindlimbs over the belt. All SCI and SCI+BDNF mice were positioned over the treadmill in the harness once a week, allowed one minute to acclimate to the moving treadmill prior to filming for two minutes. The images in Figure 2A are from video stills and had the background edited for clarity. Mice receiving daily ES were filmed twice per week on sequential days, once with stimulation off (SCI+BDNF+ES^OFF^), before daily ES sessions, and again while receiving ES the next day (SCI+BDNF+ES^ON^).

Experimental DREADD mice were injected with saline or CNO (5mg/kg) once per week, on sequential days, and filmed 40 minutes after. For each mouse, we quantified the number of ankle flexions, dorsal steps, and plantar steps performed in the 2-minute scoring window. Values from both hindlimbs were summed for subsequent comparisons. We compared these values individually and as a summed total referred to as “step events”. Additionally, for each mouse, we recorded the amount of time which mice exhibited one or more symptoms of hyperreflexia in the trunk and/or hindlimbs during the 2-minute recording on the treadmill and calculated this as a percentage to assign each mouse a weekly “hyperreflexia score.” Hindlimb stepping and hyperreflexia were calculated independently but often observed simultaneously in opposing limbs. For overground scoring, mice were placed in an open field and allowed to freely explore for 4 minutes during which we obtained scores using the Basso Mouse Scale (BMS) for post-SCI locomotor improvements.

### Spinal cord preparations

For electrophysiology experiments, 10- to 16-week-old uninjured and transected (SCI) mice were anesthetized with ketamine (150 mg/kg) and xylazine (15 mg/kg), decapitated, and eviscerated. Spinal cords were then removed in ice-cold dissecting solution containing the following (in mM): 222 glycerol, 3 KCl, 11 glucose, 25 NaHCO_3_, 1.3 MgSO_4_, 1.1 KH_2_PO_4_, and 2.5 CaCl_2_ and aerated with 95% O_2_ and 5% CO_2_ (Husch et al., 2012). The lumbar spinal cord (T11-L5) was sectioned transversely (300 µm) using a vibrating microtome (Leica Microsystems). Slices were next transferred to artificial cerebrospinal fluid containing the following (in mM): 111 NaCl, 3 KCl, 11 glucose, 25 NaHCO_3_, 1.3 MgSO_4_, 1.1 KH_2_PO_4_, and 2.5 CaCl_2_ at 37°C for 30 minutes and then passively equilibrated to room temperature for at least 1 hour before recording. Dissecting and recording solutions were continuously aerated with 95%/5% O_2_/CO_2_.

### Patch-clamp recordings

Electrophysiology experiments were performed at least 24 hours after final treadmill and ES sessions. All recordings were performed at room temperature. Fluorescently labeled (tdTomato) RORβ neurons were visualized with a 63x objective lens on a BX51WI scope (Olympus) using LED illumination (Andor Mosaic System). Patch electrodes were pulled to tip resistances of 5-8 MΩ using a multistage puller (Sutter Instruments) and were filled with intracellular solution, which contained the following (in mM): 128 K-gluconate, 10 HEPES, 0.0001 CaCl_2_, 1 glucose, 4 NaCl, 5 ATP, 0.3 GTP. Data were collected with a Multiclamp 700B amplifier (Molecular Devices) and Clampex software (pClamp9, Molecular Devices). Signals were digitized at 20 kHz and filtered at 4 kHz. Data analysis was performed with Clampfit (Molecular Devices).

### Hierarchical clustering

Agglomerative clustering analyses were performed using MATLAB algorithms (R2022b, The Mathworks Inc, Natick, MA), which do not require the specification of the number of clusters prior to initial analysis. Final measures for treadmill locomotion (ankle flexions, dorsal steps, plantar steps, and hyperreflexia score) were the inputs. We obtained standardized Z-scores for each variable and performed a principal component analysis (PCA) transformation on this dataset. The first three components explained 92.5% of the variance. Hierarchical clusters were determined by the correlated distance between observations based on the unweighted average distance (UPGMA) between clusters. This combination yielded the highest cophenetic coefficient. A dendrogram was constructed and the number of clusters was determined by natural divisions, which do not require the specification of the number of clusters prior to analysis, and evaluated by silhouette analyses. For comparative analysis, intracellular data from uninjured (N=19, n=37) and untreated SCI (N=5, n=15) mice that were not filmed, and therefore not in the cluster analysis, were included along with the data from our functionally-clustered mice. Data from the untreated SCI mice were included with data from Cluster 1. Data from uninjured mice are represented in a separate uninjured group.

### Histology

Following the final observations, animals injected with DREADD or control vectors were anesthetized and transcardially perfused with 0.1M phosphate buffered saline (PBS) followed by 4% paraformaldehyde (PFA). Spinal cords were then harvested from each animal, post-fixed in 4% PFA for 2-3 hours, and then placed in a solution of 30% sucrose in PBS for at least 24 hours to cryoprotect. Tissue was then blocked in OCT compound (Fischer Scientific International, Inc., Hampton NH) over dry ice and stored at -80°C. Spinal cords were sectioned (25µm) transversely on a frozen cryostat (Microm HM 505 E) and directly mounted onto slides in a serial fashion so that subsequent sections were sampled approximately 200 microns caudally from the previous. All slides were washed in 0.1M PBS solution before blocking with solution containing donkey serum (5%), BSA (1%), fish gelatin (1%), Triton (0.2%) and 0.1M PBS. Slides were then incubated with rabbit-anti-Pax2 (1:800, Sigma-Aldrich) primary antibody overnight at 4°C and secondary antibody (anti-rabbit-Alexa Fluor 647, 1:400, Jackson ImmunoResearch, Inc., West Grove, PA) for 3-4 hours in the dark. Slides were cover slipped using Fluoromount-G (Thermo Fisher Scientific, Inc., Waltham, MA), sealed, and stored until imaged. Images were acquired as sequential z stacks at 20x using a Leica DM6 fluorescent microscope with Thunder processing (Leica Microsystems, GmbH., Wetzlar, DE) using constant laser settings for individual slides. Images were processed offline using ImageJ software with adjustments to brightness and contrast.

### Statistical analyses

Statistical tests and *post hoc* analyses used are stated for each experiment and were performed with GraphPad Prism (Version 10.0.2, GraphPad Software, Boston, MA). All results were presented as mean ± SD, unless otherwise stated. Statistical significance was set at *p* < 0.05. Equality of group variance was determined using the Brown-Forsythe test and the distribution of the data was determined by Shapiro-Wilk normality test. Comparisons between treatments for behavioral data (SCI vs. SCI+BDNF) were performed using unpaired *t*-tests on parameters that satisfied both homogeneity of variance and normality of distribution, and using the Mann Whitney test for those that did not. For within-subject comparisons between treatment conditions (ES^OFF^ vs. ES^ON^ or saline vs. CNO), we performed two-way repeated measures ANOVA with Fisher’s LSD. Due to differences in the heteroskedasticity and distribution of the data at any given week, matched data in significant post hoc tests were also compared using Wilcoxon matched-pairs signed rank test in such instances to further confirm significant differences at the weeks identified by two-way repeated measures ANOVA analysis. Pair-wise comparisons of treadmill locomotion taken from SCI+BDNF+ES mice were exclusive to mice which had measures of treadmill performance under both conditions (ES^OFF^ vs. ES^ON^) for a given week to include experimental mice in which wires failed before intended final timepoints. Differences in behavioral and electrophysiological data between clusters were determined using one-way ANOVA with Tukey’s post hoc test for datasets that satisfied both homogeneity of variance and normality of distribution and using Welch’s ANOVA with Tamhane’s T2 post hoc for those that did not. Welch’s ANOVA is generally more robust to violations in either parameter and is the correct test in principle for such comparisons, although the same differences were identified using standard ANOVA and Tukey’s post hoc tests as well. Contingency tables were assessed to determine differences in the proportion of observed events and analyzed using two-sided Fisher’s exact test.

